# ATP-driven Non-equilibrium Activation of Kinase Clients by the Molecular Chaperone Hsp90

**DOI:** 10.1101/2020.05.10.087577

**Authors:** Huafeng Xu

## Abstract

The molecular chaperone 90-kDa heat-shock protein (Hsp90) assists the late-stage folding and activation of diverse types of protein substrates (called clients), including many kinases. Previous studies have established that the Hsp90 homodimer undergoes an ATP-driven cycle through open and closed conformations. Here I propose a model of client activation by Hsp90, which predicts that this cycle enables Hsp90 to use ATP energy to drive a client out of thermodynamic equilibrium toward its active conformation. My model assumes that an Hsp90-bound client can transition between a deactivating conformation and an activating conformation. It suggests that the cochaperone Cdc37 aids Hsp90 to activate kinase clients by differentiating between these two intermediate conformations. My model makes experimentally testable predictions, including how modulating the stepwise kinetics of the Hsp90 cycle—for example, by various cochaperones—affects the activation of different clients. My model may inform client-specific and cell-type-specific therapeutic intervention of Hsp90-mediated protein activation.

## Introduction

Many proteins depend on molecular chaperones for proper folding into their functional structures. Hsp90 is responsible for the activation (sometimes referred to as maturation) and stabilization of a wide range of client proteins—many involved in signal transduction—in the cell (1, 2), including oncogenic kinases such as v-Src (3, 4). A client of Hsp90 is often first processed by other chaperones such as Hsp70 into a partially folded state before Hsp90 completes its final maturation into the active state (5, 6). Pharmacological inhibition of Hsp90 ATPase activity results in *in vivo* degradation of its clients (7, 8), making therapeutic intervention of Hsp90 function a promising approach to the treatment of a variety of diseases caused by aberrant signal transduction, such as cancer (9), and by protein misfolding and aggregation, such as Alzheimer’s disease (10). Unraveling the mechanism by which Hsp90 activates its clients may help predict how the activities of different clients respond to different Hsp90 modulations, which may enable the rational targeting of Hsp90 and its cochaperones in order to achieve better potency and client or cell-type specificity, thus improving therapeutic efficacy and safety (9).

Highly conserved from bacteria to eukaryotes, Hsp90 functions as a homodimer. Each protomer consists of three domains: the C-terminal domain (CTD), which provides the site of dimerization, the middle domain (MD), to which clients bind, and the N-terminal domain (NTD), which harbors the ATPase capable of nucleotide-binding and ATP hydrolysis. Structural and biochemical studies have revealed that the Hsp90 dimer undergoes an ATP-driven conformational cycle (11). The nucleotide-free Hsp90 dimer preferentially adopts the open, “V”-shaped conformation (Figure 1A). The client can bind to and unbind from the open Hsp90 dimer. ATP-binding induces a reconfiguration of the nucleotide lid in NTD (Figure 1B), which exposes an additional dimerization interface and strongly biases Hsp90 toward a closed conformation where the two NTDs also dimerize (Figure 1C). This has been clearly demonstrated by electron microscopy (12) and single molecule experiments (13) in which Hsp90 incubated with an unhydrolyzable ATP analog 5’-adenylyl-β,γ-imidodiphosphate (AMPPNP) is only seen in the closed conformation. In the closed conformation, a catalytic loop containing a conserved arginine from MD is repositioned to complete the catalytic center for ATP hydrolysis. ATP hydrolysis leads to a compact ADP-bound Hsp90 dimer (Figure 1D) and perhaps the loss of the client-binding surface (12). Subsequent ADP release then returns the Hsp90 dimer to its open conformation.

**Figure 1.**
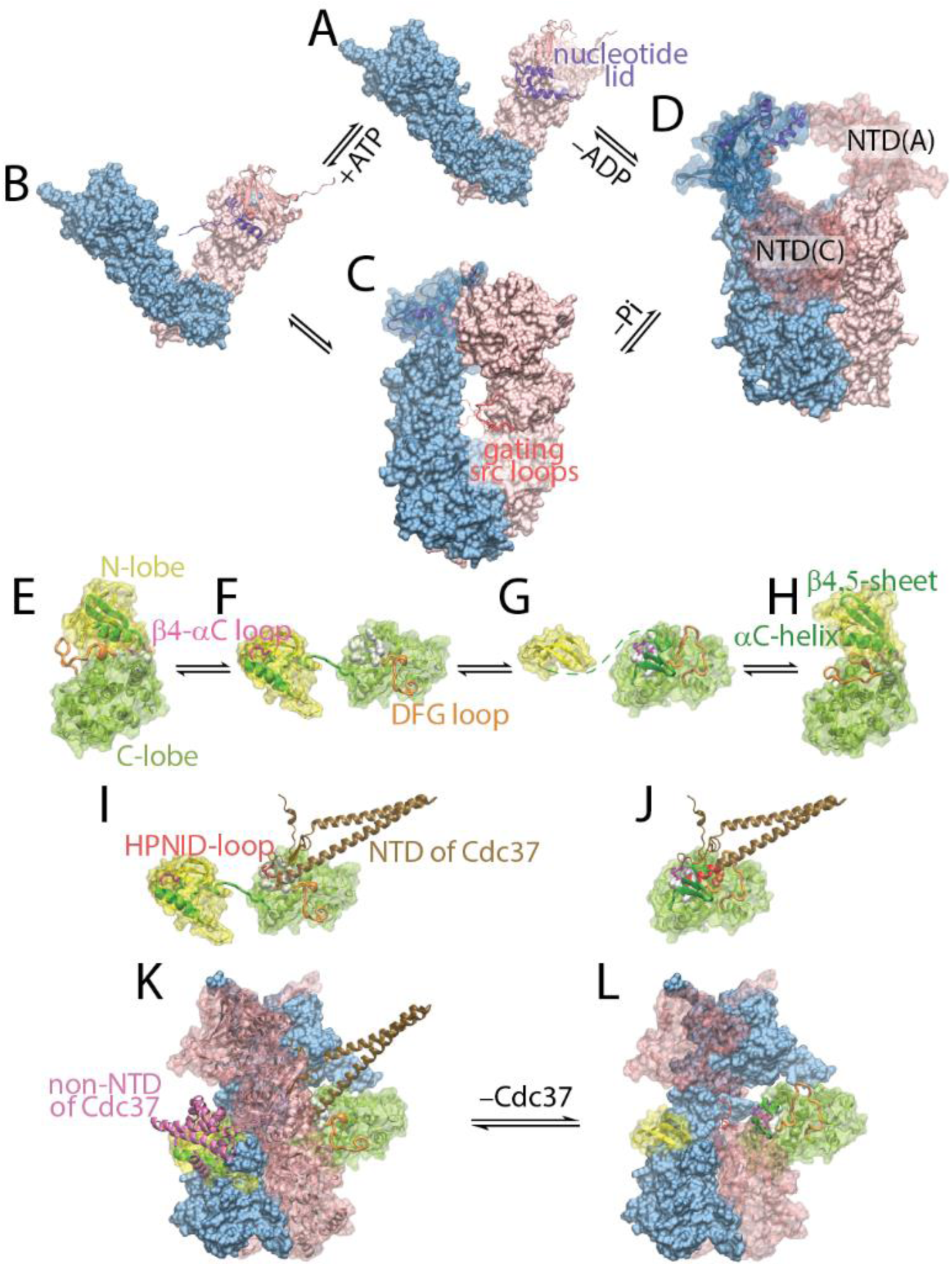
Structural basis for my model of non-equilibrium activation of client kinases by Hsp90. The underlined texts below are PDB IDs and chain IDs. The Hsp90 homodimer goes through four states in **A** to **D**. (**A**) Hsp90_open_: nucleotide-free and open; <u>2IOQ</u>. (**B**) 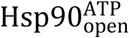: ATP-bound and open; <u>2IOQ</u> but with the NTD of one chain replaced with the ATP-bound NTD from the structure <u>2CG9</u>. ATP-binding induces a structural change in the nucleotide lid in NTD. (**C**) 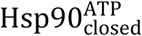: ATP-bound and closed; <u>2CG9</u>. The hole between the two MDs in the closed Hsp90 dimer is gated by a flexible, often disordered src loop (the src loops from <u>2CG9, 5FWM, 2IOP</u>, and <u>4IYN</u> are superimposed). (**D**) 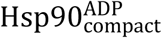: ADP-bound and compact; <u>2IOP</u>. The ADP-bound NTD may have a number of orientations with respect to MD: *e.g.*, the NTD of chain A may move to the position occupied by the NTD of chain C in the crystal structure, resulting in a compact conformation consistent with electron microscopy images (12) but clashing with the bound kinase client. In B, C, and D, the nucleotides are shown in spheres. The kinase client may transition through four states in **E** to **H**. (**E**) The inactive conformation: *i*; <u>1HCL:A</u>. (**F**) The deactivating conformation: *d*; <u>5FWM:K</u>. (**G**) The hypothetical maturing conformation: *m*. (**H**) The active conformation: *a*; <u>1FIN:A</u>. The deactivating structure is of the client kinase Cdk4 in the Hsp90-Cdc37-Cdk4 complex; the inactive and active structures are of Cdk2. (**I**) Complex between Cdc37 and Cdk4 in the *d*-state (<u>5FWM:K+E)</u>. (**J**) Steric clash prevents Cdc37 from binding to Cdk4 in the *m*-state. Three residues (in spheres) in the DFG loop—in addition to perhaps the β4-αC loop—would collide with Cdc37, whose clashing residues are highlighted in red. (**K**) The ternary complex of Hsp90-Cdc37-Cdk4 (<u>5FWM</u>), where the client kinase is in the *d*-state. (**L**) The hypothetical complex of the closed Hsp90 dimer with the client kinase in the *m*-state.

An unsettled question is how Hsp90 activates its clients through its ATP cycle (14). Compared to other ATP-driven chaperones such as chaperonins and Hsp70, Hsp90 is a very slow ATPase, with a turn-over rate of 0.05-1.2 ATP per minute and a large *K_m_* in the range of 15 to 250 µM (11). ATP hydrolysis, however, appears indispensable to Hsp90 for its in vivo function (15, 16) and client activation (17). What is the use of the spent ATP free energy? Some cochaperones aid Hsp90 in the activation of a specific class of clients, such as Cdc37 in the case of kinases (18, 19). Their roles in the client activation remain an open question. A number of cochaperones, such as Aha1 and p23, modulate Hsp90 function by either accelerating or slowing down the ATP cycle. It is puzzling why the altered cycle timing affects client activation (17).

Here I propose a model of Hsp90-mediated activation of client kinases, which predicts that Hsp90 uses the free energy from ATP hydrolysis to elevate the active fraction of a client above its Hsp90-free equilibrium value. A key assumption of my model is that the client can transition between two intermediate conformations—one favoring activation and the other deactivation— when it is clamped by Hsp90 in the closed conformation. I will discuss the structural basis of this assumption and the likely types of conformational changes, using as an example two hypothetical kinase intermediate conformations; I will also present a mathematical proof that such conformational transition in an Hsp90-bound client is necessary for its non-equilibrium activation. My model suggests a new thermodynamic criterion that distinguishes between client and non-client proteins; it provides a mechanism by which Cdc37 expands Hsp90’s kinase clientele. It explains how cochaperones such as Aha1 affect client activation by changing the kinetic rates of different steps in Hsp90’s ATP cycle. My model implies that Hsp90 is not only involved in the folding of nascent protein chains but also in the maintenance of their activity. This and other predictions of my model subject it to experimental tests.

## Results

Theoretical (20) and simulation studies (21) have suggested that proteins such as kinases go through locally unfolded intermediate conformations in order to facilitate the inactive-to-active transition. Hsp90 binds to the kinase clients in these intermediate conformations, as revealed by the structures of the Hsp90-Cdc37-Cdk4 complex (22), including an atomic structure by cryoEM at 3.9 Å resolution (23). In this structure, the closed Hsp90 dimer clamps its client kinase, which adopts a bilobial structure with its N- and C-lobes separated to two sides of the clamp (Figure 1K). The two individual lobes are largely folded, but their interface is completely broken (Figure 1F), which is consistent with the observation that Hsp90 is mostly involved in the final stage of protein folding and activation. Evidence has emerged that instead of adopting a single structure, a client bound to Hsp90 is in a fluid conformational ensemble (24).

It is reasonable to presume that the intermediate conformations can be divided into two categories based on whether they are more likely to transition into the active state or into the inactive state. Here I assume in a simple model that the client can adopt two locally unfolded intermediate states (Figure 1F, G; I will discuss later these hypothetical kinase structures in detail): in the deactivating state (the *d*-state), the client is strongly disposed to fold into the inactive state (the *i*-state); in the maturing state (the *m*-state), the client is poised to fold into the active state (the *a*-state). The client can transition between the *i*- and the *d*-states, between the *d*- and the *m*-states, and between the *m*- and the *a*-states (Figure 1E-H). Hsp90 can bind to the client in the intermediate states but not in the inactive or active states.

Stable binding between the client and the open Hsp90 dimer will likely require extensive contacts between the client and Hsp90 and thus hinder the *d* ⇌ *m* conformational transition. In the closed Hsp90 dimer, however, the client is clamped between the two Hsp90 protomers and it cannot move away even if it break its contacts with Hsp90, thus allowing it to interconvert between the *d*- and the *m*-states while still bound to Hsp90. The structures of the closed Hsp90 dimer—conserved across great evolutionary distances—indeed hint at such conformational transitions. A large hole exists at the center of the dimer, but it is gated by two flexible src loops (Figure 1C). These gating loops prevent folded domains from slipping through the hole and thus keep the client trapped, but the remaining space appears large enough to permit unfolded amino acid chains to thread through.

I thus propose a model (Figure 2 and Table 1) that couples Hsp90’s ATP-driven conformational cycle with the client’s conformational changes. In my model, Hsp90 cycles through 4 states: 1) the open and nucleotide-free state (Hsp90_open_), which interconverts—by ATP binding—with 2) the open and ATP-bound state 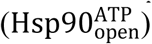, which overwhelmingly converts to the more stable 3) closed and ATP-bound state 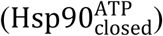, which, following ATP hydrolysis, converts to 4) the compact and ADP-bound state 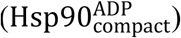, which, upon ADP release, returns to the open and nucleotide-free state. Although each state consists of multiple conformational sub-states (25), and the two protomers hydrolyze ATP at a deterministic sequential order (26), for simplicity these details are omitted from the current model. The client in the *d*- or the *m*-state, using hydrophobic patches exposed by the local unfolding, can reversibly bind to Hsp90_open_ and 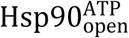. When the client is in the solution or when it is clamped in 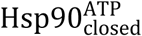, it can interconvert between the *d*- and *m*-states.

**Table 1.** Reactions and kinetic parameters for Hsp90-activation of v-Src kinase. Hsp90 is abbreviated as H90, and Cdc37 as C37. The boldfaced values are fitted to the experimental data of Hsp90-activation of v-Src. The underlined values are uniquely determined from other parameters so that the total free energy changes in any reaction cycle is zero. The kinetic parameters of the Hsp90 cycle (reactions 1 to 4) are derived from kinetic analysis of nucleotide-driven changes in fluorescence resonance energy transfer in labeled Hsp90 (25), with the *k_f_* of reaction 1 adjusted by a factor of 2 so that the ATPase has a *K_m_* of 250 µM (11). The kinetic parameters for v-Src conformational transitions—experimentally unavailable—are chosen arbitrarily, except for *k_f_* of reaction 7. The *k_f_* of reactions 11d and 11m reflects a 20-fold acceleration of the closing of the Hsp90 dimer induced by the client (33, 34). Other than this acceleration, the client is assumed not to affect the kinetics of other steps of the ATP-driven cycle (reactions 8 to 13). I also assume that the rates of client’s conformational transition is the same in the closed Hsp90 dimer (reaction 14) as in the solution (reaction 6). All protein-protein association rate constants are taken to be 1 µM^−1^s^−1^, a reasonable value, except for reactions 19a and 20, which should be slower because Cdc37 has to adopt a specific conformation in order to simultaneously bind to both Hsp90 and the client. The dissociation rate constant of Cdc37 from Hsp90 (reaction 15) is computed from its binding constant *K_D_* = 1.4 µM (43, 44). The dissociation rate constant of reaction 20 is arbitrary because the model predictions are insensitive to its value.

| No. | Reaction | $k_f$ | $k_r$ |
| --- | --- | --- | --- |
| 1 | $\text{H90}_{\text{open}} + \text{ATP} \rightleftharpoons \text{H90}_{\text{open}}^{\text{ATP}}$ | $6.7 \times 10^{-4} \mu\text{M}^{-1}\text{s}^{-1}$ | $1.45 \times 10^{-1} \text{s}^{-1}$ |
| 2 | $\text{H90}_{\text{open}}^{\text{ATP}} \rightleftharpoons \text{H90}_{\text{closed}}^{\text{ATP}}$ | $1.7 \times 10^{-2} \text{s}^{-1}$ | $5 \times 10^{-5} \text{s}^{-1}$ |
| 3 | $\text{H90}_{\text{closed}}^{\text{ATP}} \rightleftharpoons \text{H90}_{\text{compact}}^{\text{ADP}} + \text{Pi}$ | $3.3 \times 10^{-1} \text{s}^{-1}$ | <u>$9.3 \times 10^{-14} \mu\text{M}^{-1}\text{s}^{-1}$</u> |
| 4 | $\text{H90}_{\text{compact}}^{\text{ADP}} \rightleftharpoons \text{H90}_{\text{open}} + \text{ADP}$ | $1 \text{s}^{-1}$ | $1 \times 10^{-1} \mu\text{M}^{-1}\text{s}^{-1}$ |
| 5 | $i \rightleftharpoons d$ | $1 \text{s}^{-1}$ | $5 \times 10^2 \text{s}^{-1}$ |
| 6 | $d \rightleftharpoons m$ | $1 \times 10^{-1} \text{s}^{-1}$ | $1 \times 10^{-1} \text{s}^{-1}$ |
| 7 | $m \rightleftharpoons a$ | **$6.5 \times 10^1 \text{s}^{-1}$** | $1 \text{s}^{-1}$ |
| 8d | $d + \text{H90}_{\text{open}} \rightleftharpoons d \cdot \text{H90}_{\text{open}}$ | $1 \mu\text{M}^{-1}\text{s}^{-1}$ | **$5.6 \times 10^{-3} \text{s}^{-1}$** |
| 8m | $m + \text{H90}_{\text{open}} \rightleftharpoons m \cdot \text{H90}_{\text{open}}$ | $1 \mu\text{M}^{-1}\text{s}^{-1}$ | **$6.2 \times 10^{-1} \text{s}^{-1}$** |
| 9d | $d + \text{H90}_{\text{open}}^{\text{ATP}} \rightleftharpoons d \cdot \text{H90}_{\text{open}}^{\text{ATP}}$ | Same as 8d | Same as 8d |
| 9m | $m + \text{H90}_{\text{open}}^{\text{ATP}} \rightleftharpoons m \cdot \text{H90}_{\text{open}}^{\text{ATP}}$ | Same as 8m | Same as 8m |
| 10d | $d \cdot \text{H90}_{\text{open}} + \text{ATP} \rightleftharpoons d \cdot \text{H90}_{\text{open}}^{\text{ATP}}$ | Same as 1 | Same as 1 |
| 10m | $m \cdot \text{H90}_{\text{open}} + \text{ATP} \rightleftharpoons m \cdot \text{H90}_{\text{open}}^{\text{ATP}}$ | Same as 1 | Same as 1 |
| 11d | $d \cdot \text{H90}_{\text{open}}^{\text{ATP}} \rightleftharpoons d \cdot \text{H90}_{\text{closed}}^{\text{ATP}}$ | $3.3 \times 10^{-1} \text{s}^{-1}$ | **$5.4 \times 10^{-3} \text{s}^{-1}$** |
| 11m | $m \cdot \text{H90}_{\text{open}}^{\text{ATP}} \rightleftharpoons m \cdot \text{H90}_{\text{closed}}^{\text{ATP}}$ | Same as 11d | Same as 2 |
| 12d | $d \cdot \text{H90}_{\text{closed}}^{\text{ATP}} \rightleftharpoons d \cdot \text{H90}_{\text{compact}}^{\text{ADP}} + \text{Pi}$ | Same as 3 | Same as 3 |
| 12m | $m \cdot \text{H90}_{\text{closed}}^{\text{ATP}} \rightleftharpoons m \cdot \text{H90}_{\text{compact}}^{\text{ADP}} + \text{Pi}$ | Same as 3 | Same as 3 |
| 13d | $d \cdot \text{H90}_{\text{compact}}^{\text{ADP}} \rightleftharpoons d \cdot \text{H90}_{\text{open}} + \text{ADP}$ | Same as 4 | $1.8 \times 10^{-2} \mu\text{M}^{-1}\text{s}^{-1}$ |
| 13m | $m \cdot \text{H90}_{\text{compact}}^{\text{ADP}} \rightleftharpoons m \cdot \text{H90}_{\text{open}} + \text{ADP}$ | Same as 4 | $2.0 \mu\text{M}^{-1}\text{s}^{-1}$ |
| 14 | $d \cdot \text{H90}_{\text{closed}}^{\text{ATP}} \rightleftharpoons m \cdot \text{H90}_{\text{closed}}^{\text{ATP}}$ | Same as 6 | Same as 6 |
| 15 | $\text{C37} + \text{H90}_{\text{open}} \rightleftharpoons \text{C37} \cdot \text{H90}_{\text{open}}$ | $1 \mu\text{M}^{-1}\text{s}^{-1}$ | $1.4 \text{s}^{-1}$ |
| 16 | $\text{C37} + d \rightleftharpoons d \cdot \text{C37}$ | $1 \mu\text{M}^{-1}\text{s}^{-1}$ | $5.1 \times 10^{-3} \text{s}^{-1}$ |
| 17 | $d + \text{C37} \cdot \text{H90}_{\text{open}} \rightleftharpoons d \cdot \text{C37} \cdot \text{H90}_{\text{open}}$ | Same as 16 | $4.8 \times 10^{-4} \text{s}^{-1}$ |
| 18a | $d \cdot \text{C37} + \text{H90}_{\text{open}} \rightleftharpoons d \cdot \text{C37} \cdot \text{H90}_{\text{open}}$ | $1 \mu\text{M}^{-1}\text{s}^{-1}$ | $1.3 \times 10^{-1} \text{s}^{-1}$ |
| 18n | $d \cdot \text{C37} + \text{H90}_{\text{open}}^{\text{ATP}} \rightleftharpoons d \cdot \text{C37} \cdot \text{H90}_{\text{open}}^{\text{ATP}}$ | Same as 18a | $2.8 \times 10^{-2} \text{s}^{-1}$ |
| 19a | $\text{C37} + d \cdot \text{H90}_{\text{open}} \rightleftharpoons d \cdot \text{C37} \cdot \text{H90}_{\text{open}}$ | $2 \times 10^{-1} \mu\text{M}^{-1}\text{s}^{-1}$ | $2.4 \times 10^{-2} \text{s}^{-1}$ |
| 19n | $\text{C37} + d \cdot \text{H90}_{\text{open}}^{\text{ATP}} \rightleftharpoons d \cdot \text{C37} \cdot \text{H90}_{\text{open}}^{\text{ATP}}$ | $1 \mu\text{M}^{-1}\text{s}^{-1}$ | $2.5 \times 10^{-2} \text{s}^{-1}$ |
| 20 | $\text{C37} + d \cdot \text{H90}_{\text{closed}}^{\text{ATP}} \rightleftharpoons d \cdot \text{C37} \cdot \text{H90}_{\text{closed}}^{\text{ATP}}$ | $1 \times 10^{-1} \mu\text{M}^{-1}\text{s}^{-1}$ | $1 \times 10^{-1} \text{s}^{-1}$ |
| 21 | $d \cdot \text{C37} \cdot \text{H90}_{\text{open}} + \text{ATP} \rightleftharpoons d \cdot \text{C37} \cdot \text{H90}_{\text{open}}^{\text{ATP}}$ | Same as 1 | $3 \times 10^{-2} \text{s}^{-1}$ |
| 22 | $d \cdot \text{C37} \cdot \text{H90}_{\text{open}}^{\text{ATP}} \rightleftharpoons d \cdot \text{C37} \cdot \text{H90}_{\text{closed}}^{\text{ATP}}$ | Same as 11d | $2.2 \times 10^{-1} \text{s}^{-1}$ |

**Figure 2.**
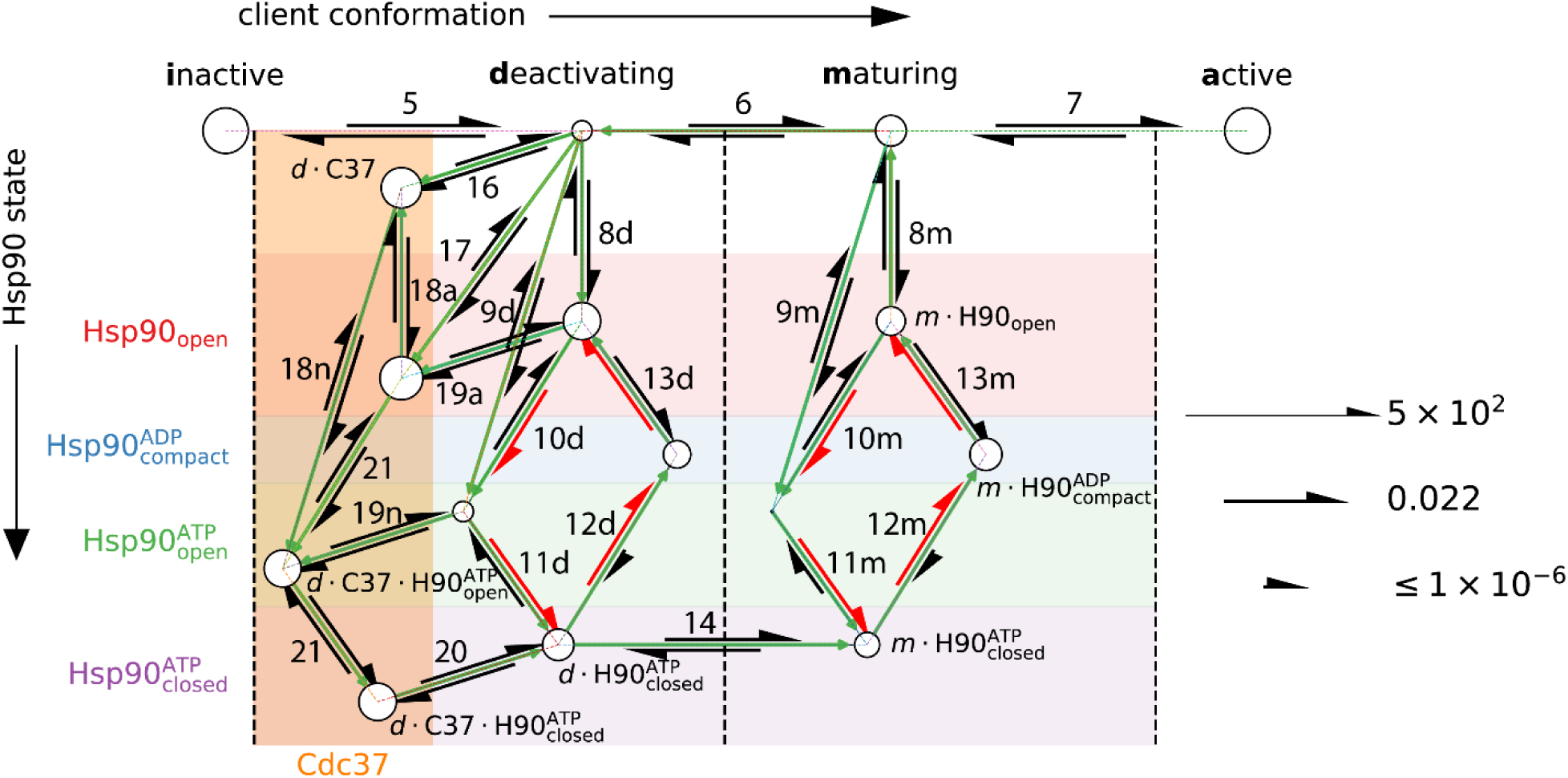
The reactions in Hsp90-mediated non-equilibrium activation of v-Src and the reactive fluxes at the steady state. Each circle represents a molecular species; the radius of the circle is linear in the logarithm of its steady state concentration. The top four circles represent the free client molecule in its four different conformations; the others represent complexes between the client and Hsp90. The molecular species are divided by the vertical dashed lines according to the client conformation, and into the colored horizontal bands according to the conformational and nucleotide state of Hsp90. The ones that lie in the orange vertical band are ternary complexes of Hsp90-client-Cdc37. The Hsp90 ATP cycles are highlighted in red. The intermediate molecular species in the Cdc37-assisted activation pathway are labeled. The directions of the steady state reactive fluxes are indicated by the green arrows. The reactions are labeled with the reaction numbers in Table 1. The reaction conditions are [Hsp90] = [Cdc37] = 1.3 µM, [v-Src] = 0.32 µM, [ATP] = 20 µM, and [ADP] = [Pi] = 1 µM.

My model implies the following pathway of client activation. First, an inactive client locally unfolds to the *d*-state and binds to an open Hsp90 dimer, which closes upon ATP binding. The client then transitions from the *d*- to the *m*-state. Hsp90 hydrolyzes ATP. After ADP-release, the Hsp90 dimer reopens, allowing the maturing client to unbind and fold into its active conformation. Denoting a complex between molecules *x* and *y* as *x* · *y*, this *d*-to-*m* pathway can be written as

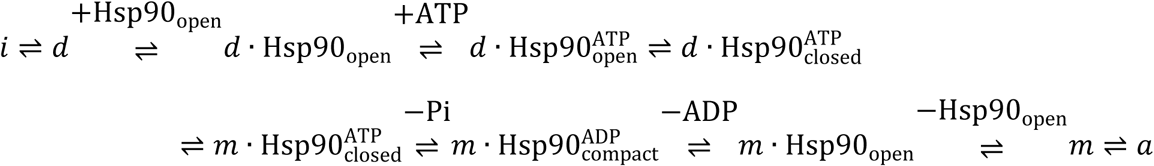

There is a corresponding *m*-to-*d* pathway—swapping *i* with *a* and *d* with *m*—that deactivates the client:

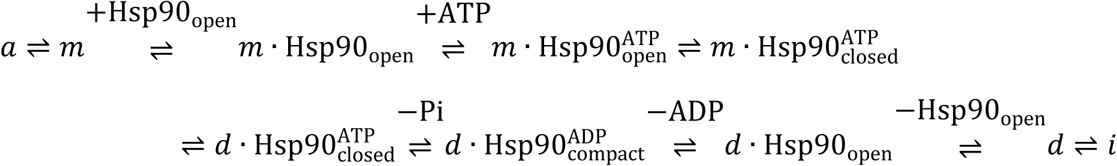

Note the directionality of the ATP-driven Hsp90 conformational cycle (one ATP molecule is hydrolyzed in each pathway above); the second *m*-to-*d* pathway is not the reverse of the first *d*- to-*m* pathway.

Hsp90 can drive the client toward the active state if the *d*-to-*m* pathway occurs more than the *m*- to-*d* pathway. How can this happen at the steady state? One possible mechanism is that the open Hsp90 dimer has a higher affinity for the client in the *d*-state than for the client in the *m*-state. This promotes the *d*-to-*m* pathway over the *m*-to-*d* pathway in the steps of client binding and release: a client molecule in the *d-*state is more likely than one in the *m*-state to bind to Hsp90; a client molecule in the *m*-state is more likely than one in the *d*-state to unbind from Hsp90. But it also means that the complex *d* · Hsp90_open_ is more stable than the complex *m* · Hsp90_open_ and the external free energy from ATP is needed to drive the *d*-to-*m* pathway from *d*. Hsp90_open_ to *m* · Hsp90_open_. Key to this mechanism is the overwhelming tendency of the ATP-bound Hsp90 dimer to close: 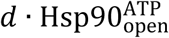 is more stable than 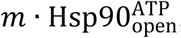, which implies a higher free energy penalty for the step of 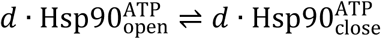 in the *d*-to-*m* pathway than for 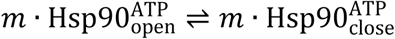 in the *m*-to-*d* pathway, but this difference is negligible in comparison to the large thermodynamic motive for the Hsp90 dimer to close in both complexes.

To test whether this mechanism is consistent with experimentally observed client activation by Hsp90, I used my model to analyze the activation of v-Src, a stringent Hsp90 client (4). The normalized activity of v-Src—measured experimentally as the activity ratio between v-Src incubated with and without Hsp90—is computed as the ratio of the active (*i.e. a*-state) concentration at the steady state to that at the Hsp90-free equilibrium (Figure 3A). The calculated normalized activity at the steady state is greater than 1 in the presence of Hsp90, affirming that the proposed mechanism enables Hsp90 to convert ATP energy into non-equilibrium stabilization of the active state of v-Src.

**Figure 3.**
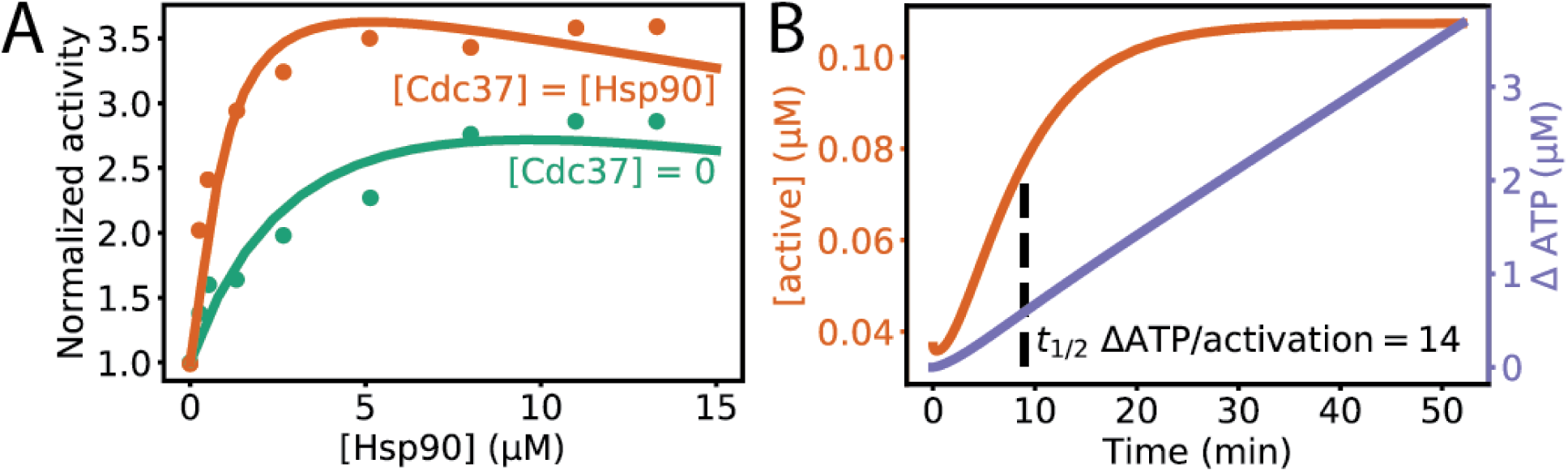
Activation of v-Src by Hsp90. (**A**) Normalized v-Src activity at the steady state at different Hsp90 and Cdc37 concentrations. The experimental data (4) are shown as points, and the results calculated from my model are shown as solid lines. (**B**) The time course of v-Src activation by Hsp90 in the presence of Cdc37 and the concomitant ATP consumption.

The more Hsp90 prefers the binding to a protein substrate in its *d*-state to the binding to the protein in its *m*-state, the more Hsp90 elevates the substrate’s steady state activity (Figure 4A). Presence of such an affinity difference may be a thermodynamic criterion that distinguishes the clients from the non-clients. But in the absence of a structural basis that systematically biases Hsp90 to bind a protein in its *d*-state stronger than the protein in its *m*-state, it is unlikely that Hsp90 solely depends on this difference in recognizing clients.

**Figure 4.**
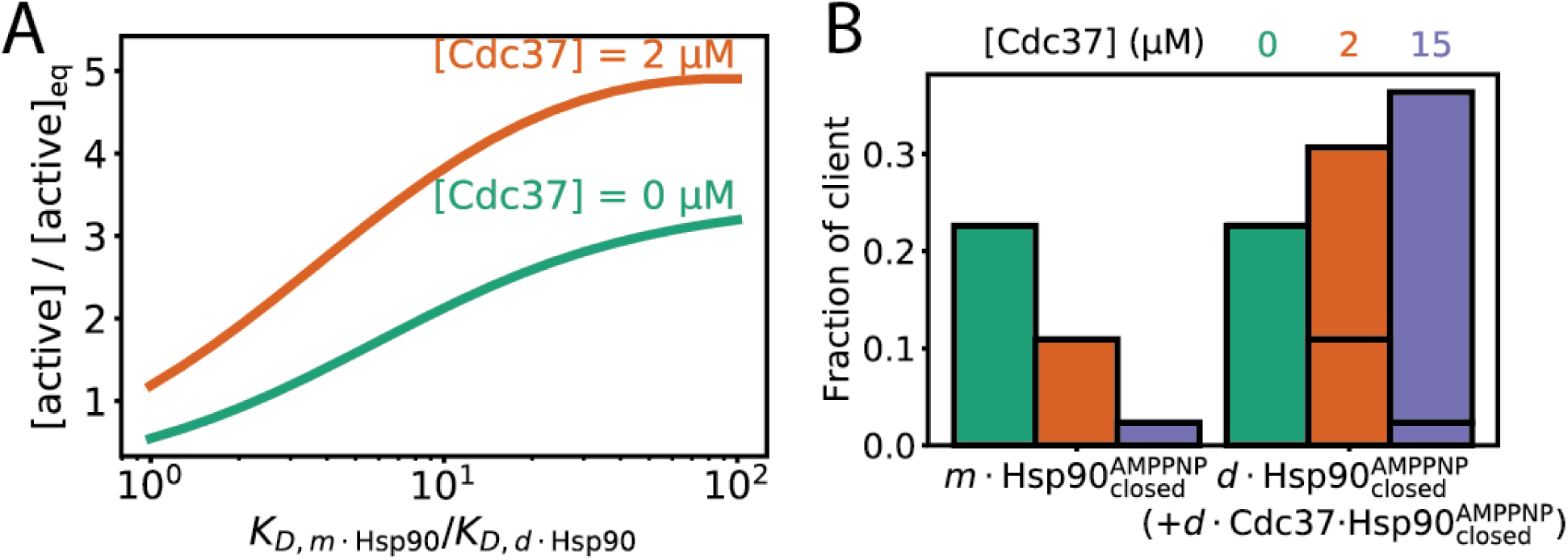
Cdc37 expands Hsp90’s clientele by recruiting deactivating kinases. (**A**) Kinase activation by Hsp90 at different ratios of its binding affinity for the *d*-state to that for the *m*-state of the kinase. Here, the *K_D_* ratio is varied while keeping the conformational equilibrium constants of the apo and the Hsp90-bound client unchanged (reactions 6 and 14 in Table 1), which is satisfied by setting *K*_*d*,closure_/*K*_*m*,closure_ = *K*_*D,m*·Hsp90_/*K*_*D,d*·Hsp90_, where *K*_*s*,closure_ is the equilibrium constant of the reaction 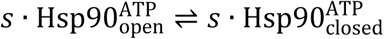 for *s* = *d, m*. In the presence of Cdc37, kinases with low *K_D_* ratios are activated by Hsp90. The reaction conditions differ from those in Figure 2 only in [Cdc37] and in that [ATP] = 250 µM. (**B**) The fraction of v-Src trapped in its *d*- and *m*-states by the closed Hsp90 dimer with the nonhydrolysable ATP analog AMPPNP at different Cdc37 concentrations. The reaction conditions are [Hsp90] = 15 µM, [v-Src] = 0.32 µM, and [AMPPNP] = 250 µM.

My model suggests a mechanism for Hsp90 to expand its clientele by cooperating with client-class-specific cochaperones (14, 27). One such example is Cdc37, which plays a critical role in activating Hsp90-dependent kinases (18). Experimental evidence (19) suggests that Cdc37 binds to a locally unfolded conformation of the client kinase and stabilizes it for Hsp90-binding. Cdc37 can form ternary complex with the locally unfolded client kinase and Hsp90 in the open and closed states (18), but it may be expelled from the complex when Hsp90 changes to the more compact conformation upon ADP hydrolysis. If Cdc37 selectively binds to the client in the *d*-state but not to the client in the *m*-state (the structural basis for this selectivity is discussed below), it will provide an additional pathway for the client to move from the *d*-state to the *m*-state through the Hsp90 conformational cycle:

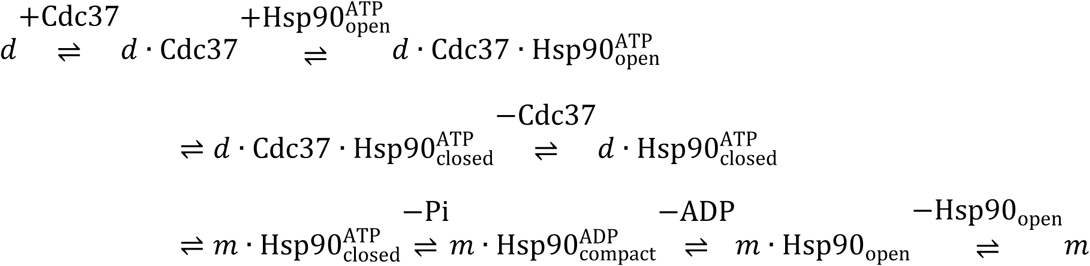

In this pathway, Cdc37 unbinds from the client in the *d*-state before the Hsp90 dimer enters the compact conformation (the unbinding can happen either before or after the ATP hydrolysis), allowing the client to transition from the *d*-state to the *m*-state in the closed Hsp90 dimer. Cdc37 thus selects the client molecules in the *d*-state to enter the Hsp90 cycle but allows some of them to exit the cycle in the *m*-state, which boosts the *d*-to-*m* transition.

My model indeed captures Cdc37’s augmentative effect on Hsp90’s non-equilibrium activation of kinases (Figures 3 and 4A). Cdc37 substantially increases the maximum normalized activity at the steady state; it also decreases the Hsp90 concentration required to achieve the maximum activation. The net reactive fluxes at the steady state (Figure 2) demonstrate that the client in the *d*-state enters the Hsp90 cycle either alone or in complex with Cdc37. There is a net flux for the client to convert from the *d*- to the *m*-state when clamped in the closed Hsp90 dimer, signifying that the *d*-to-*m* pathway dominates the *m*-to-*d* pathway. Cdc37 enables Hsp90 to elevate the steady state activity of kinases even when Hsp90 on its own cannot differentiate between the *d*- and the *m*-states (Figure 4A).

Is there a structural basis for Cdc37 to differentiate between the *d*- and the *m*-states of a client kinase? Kinase activation invariably entails the reconfiguration of its DFG activation loop from a disordered ensemble of structures into a well-ordered structure bridging the N- and C-lobes of the kinase (Figure 1E, H)(28, 29). The DFG loop of Cdk4 in the Hsp90-Cdc37-Cdk4 structure is similar to that of an inactive kinase. In contrast, superimposing the C-lobe of the active Cdk2 kinase (a close homolog of Cdk4) onto Cdk4 in the same structure results in substantial steric clashes between the DFG loop and Cdc37 (Figure 1J). These observations suggest a pair of possible structural models for the *d*- and *m*-states of a client kinase: both have the N- and C-lobes separated; the *d*-state structure has a disordered DFG loop, but the *m*-state structure has the DFG loop in its active configuration, which prevents Cdc37 binding. In addition, the β4-αC loop may move from the N-lobe side in the *d*-state to dock onto the C-lobe in the *m*-state, which would also disrupt Cdc37 binding, because Cdc37 binds to the C-lobe using a 5-residue (20HPNID24) loop mimicking the β4-αC loop; such a move seems permissible by the gated central hole in the closed Hsp90 dimer. Cdc37 can thus discriminate the *d*- and the *m*-states by the positions of the DFG-loop and perhaps of the β4-αC loop, a mechanism likely applicable across all kinase clients.

Does a client indeed change conformations while bound to Hsp90? Much remains to be learned about the structures of clients in complex with Hsp90 (30), but the molten globule state observed for a Hsp90-bound client (24) and the gated hole in the closed Hsp90 dimer structure hint at the proposed conformational change. To shore up the proposed mechanism, here I prove mathematically that for Hsp90 to use ATP energy for non-equilibrium activation of the client, the client must undergo a conformational transition while it is bound to Hsp90. This follows from a general theorem that I prove in Methods: at the steady state, two molecular species connected by a chemical reaction can only be out-of-equilibrium, *i.e.* a non-zero flux is sustained in the reaction, if this reaction is part of a reaction cycle—defined as a set of reactions that together return all molecular species to their original states (see Methods)—in which external chemical energy is consumed. If the client does not undergo a transition between two interconverting conformations, say *x* and *y*, during the time that it is bound to Hsp90, the reaction *x* ⇌ *y* cannot be part of an energy-consuming reaction cycle involving the client, Hsp90, and their complexes, because it is the only reaction in which *x* and *y* appear on the different sides of reactants and products. The steady state reactive flux between any two conformations of the apo client is thus zero, their concentration ratio is the same as the equilibrium value, and non-equilibrium activation is impossible. If non-equilibrium client activation is experimentally demonstrated (see Discussion), this theorem implies that a conformational transition must take place in the Hsp90-bound client.

I used my model to estimate ATP consumption by Hsp90 in non-equilibrium client activation. Figure 3B shows the calculated time course of the activation of v-Src—starting from the equilibrium concentrations—and the accompanying ATP consumption. In the initial phase of activation (ending when the active concentration reaches midway between the starting and the steady state values), roughly 14 ATP molecules are hydrolyzed per each activated v-Src molecule. This is considerably smaller than the ATP consumption in Hsp70-mediated folding (>100 ATP molecules per folded molecule of protein substrate)(31).

My model helps to unravel how cochaperones affect Hsp90 function by altering the kinetics of different steps in the ATP cycle. To maintain non-equilibrium activation, the cycle time has to be sufficiently fast to outpace the client’s return to equilibrium. On the other hand, Hsp90 has to dwell sufficiently long both in the open state to accommodate client binding and release and in the closed, pre-hydrolysis state to allow client conformational transition. Calculations by my model demonstrate that changes in the kinetic rates of different steps in the Hsp90 cycle affect client activation differently (Figure 5A-C).

**Figure 5.**
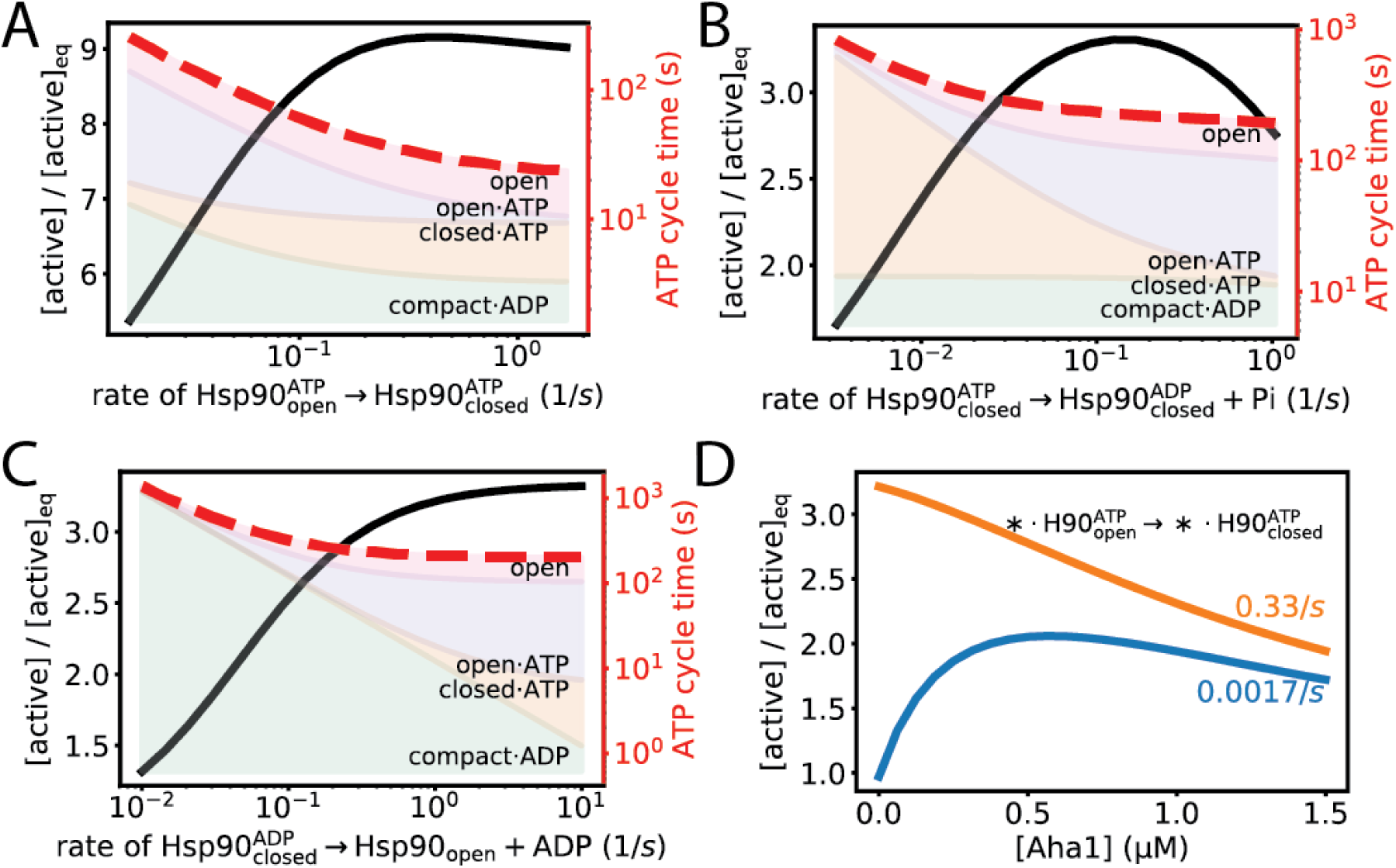
Changing the kinetic rates of the steps in the ATP cycle affects client activation by Hsp90. Normalized activity (black line and left y-axis) and ATP cycle time (red dashed line and right y axis) are calculated for different rates of (**A**) Hsp90 transition from the open to the closed conformation, (**B**) ATP hydrolysis after Hsp90 adopts the closed conformation, and (**C**) ADP release and the return of Hsp90 to the open conformation. The proportions of time spent in the four Hsp90 states—open and nucleotide-free, open and ATP-bound, closed and ATP-bound, and compact and ADP-bound—are shown as colored bands. Note the logarithmic scale of the times: for the same apparent thickness, a higher band represents a much larger occupancy than a lower band; in particular, the nucleotide-free open state has a higher occupancy than it seems. (**D**) Modulation of client activation by the cochaperone Aha1. Aha1 can augment or diminish client activation depending on whether the client itself accelerates (orange) or slows down (blue) the rate that the Hsp90 dimer closes. Here [ATP] = 250 µM, [Cdc37] = 0 µM, [Hsp90] = 1.3 µM, [client] = 0.32 µM, and [ADP] = [Pi] = 1 µM.

By considering a cochaperone’s effects on the kinetics of the cycle, my model can predict its role in the activation of different clients by Hsp90. For instance, the cochaperone Aha1 stimulates the ATP turn-over by accelerating the change from the open to the closed Hsp90 dimer, but it likely stabilizes a conformation that is not fully hydrolysis-competent (32) and it may have to unbind from Hsp90 before ATP hydrolysis can proceed. It thus have two competing effects: 1) accelerating the conformational closure of the Hsp90 dimer, which enhances client activation (Figure 5A); 2) slowing down the step of ATP hydrolysis, which may compromise client activation (Figure 5B). To analyze Aha1’s overall effect on client activation, I included Aha1 into my model (Table 2), and calculated the normalized activities of two hypothetical clients at different Aha1 concentrations (Figure 5D). Different clients either accelerate (33, 34) or slow down (35) the cycle. For a client that on its own slows down the Hsp90 dimer closure, Aha1 can augment its steady state activity by accelerating the closure. For a client that on its own accelerates the closure, however, Aha1 may diminish its steady state activity by slowing down the ATP hydrolysis. This analysis may explain the paradoxical observation that Aha1 downregulation promotes the Hsp90-dependent folding of a common disease variant of cystic fibrosis transmembrane conductance regulator (36).

**Table 2.** Reactions and kinetic parameters for the cochaperone Aha1. The dissociation rate constants are computed from the binding affinities measured by surface plasmon resonance (32). I assume that Aha1 accelerates the closure of the Hsp90 dimer by 10-fold (45). The rate of client-bound Hsp90 dimer closure in the presence of Aha1 (reactions 26d, 26m) is given by the larger *k_f_* of the reaction 11d and of the reaction 25; the reverse rate is computed from the reaction cycle of the indicated reactions.

| No. | Reaction | $k_f$ | $k_r$ |
| --- | --- | --- | --- |
| 23a | $\text{H90}_{\text{open}} + \text{Aha1} \rightleftharpoons \text{Aha1} \cdot \text{H90}_{\text{open}}$ | $1 \mu\text{M}^{-1}\text{s}^{-1}$ | $1.2 \text{s}^{-1}$ |
| 23n | $\text{H90}_{\text{open}}^{\text{ATP}} + \text{Aha1} \rightleftharpoons \text{Aha1} \cdot \text{H90}_{\text{open}}^{\text{ATP}}$ | Same as 1a | Same as 1a |
| 24 | $\text{H90}_{\text{closed}}^{\text{ATP}} + \text{Aha1} \rightleftharpoons \text{Aha1} \cdot \text{H90}_{\text{closed}}^{\text{ATP}}$ | $1 \mu\text{M}^{-1}\text{s}^{-1}$ | $1.6 \times 10^{-1} \text{s}^{-1}$ |
| 25 | $\text{Aha1} \cdot \text{H90}_{\text{open}}^{\text{ATP}} \rightleftharpoons \text{Aha1} \cdot \text{H90}_{\text{closed}}^{\text{ATP}}$ | $1.7 \times 10^{-1} \text{s}^{-1}$ | $6.67 \times 10^{-5} \text{s}^{-1}$ |
| 26d | $d \cdot \text{Aha1} \cdot \text{H90}_{\text{open}}^{\text{ATP}} \rightleftharpoons d \cdot \text{Aha1} \cdot \text{H90}_{\text{closed}}^{\text{ATP}}$ | Max(25 and 11d) | From 11d, 23n, 24 |
| 26m | $m \cdot \text{Aha1} \cdot \text{H90}_{\text{open}}^{\text{ATP}} \rightleftharpoons m \cdot \text{Aha1} \cdot \text{H90}_{\text{closed}}^{\text{ATP}}$ | Max(25 and 11m) | From 11m, 23n, 24 |

## Discussion

My model predicts that Hsp90, like two other ATP-consuming chaperones Hsp70 (31) and GroEL (37), is an agent of non-equilibrium protein-folding. It suggests that Hsp90 is not only involved in the initial folding of nascent protein chains, but it is also responsible for maintaining the activity of mature clients, which requires sustained ATP expenditure. This prediction is consistent with Hsp90’s role at buffering phenotypic variations against destabilizing mutations to client proteins (38) and with the experimental observation that Hsp90 ATPase inhibitors such as geldanamycin can suppress client protein activities and induce their degradation in cells (39). Given that geldanamycin’s effect *in vivo* may be more apparent on the nascent chains than on the mature proteins (40, 41), it is worthwhile to directly test this prediction by an *in vitro* experiment: first measure the activity of a client kinase maintained in the steady state with Hsp90 and an ATP regeneration system, then add geldanamycin to stop the Hsp90 cycle and measure the kinase activity after a new steady state is established. A decrease in activity will demonstrate ATP-driven non-equilibrium client activation by Hsp90.

The key assumption of my model is the existence of two interconverting intermediate client conformations, one favoring activation and the other deactivation. Here I propose the following experiment to test this assumption. A client kinase is first trapped in the closed Hsp90 dimer by incubation with a nonhydrolysable ATP analog such as AMPPNP. An ATP competitive inhibitor, perhaps geldanamycin, is then added in excess, opening the Hsp90 dimer and releasing the client, and the activity of the client is measured. My model predicts that at increasing concentrations of Cdc37 during the incubation, higher fractions of the trapped client will be in the *d*-state (Figure 4B) and forming the ternary complex 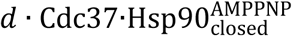, leaving smaller fractions in the *m*-state for ready activation. Consequently, in the initial moments after the client release, the measured activity will increase more slowly at higher Cdc37 concentrations.

My model predicts that client activation by Hsp90 may be promoted or suppressed by modulating the stepwise kinetics of the ATP cycle. These predictions can be tested by designing Hsp90 mutants that alter the stability of different conformational states of Hsp90 and testing their effects on the timing of the ATP cycle and on client activation (17). My model implies that the same cochaperone may affect different clients differently, depending on how the cochaperone and how the client each alters the timings of the ATP cycle, which may be tested experimentally. In addition, my model implies that the occupancies of Hsp90 conformational states will change with the concentration levels of different clients and of various cochaperones, which may help explain the observation that 17-allylaminogeldanamycin, a clinical-stage Hsp90 inhibitor, binds to Hsp90 in tumor cells with about two-orders-of-magnitude higher affinity than to Hsp90 in normal cells (42). These implications of my model raise the possibility of client-specific or cell-type-specific targeting of Hsp90 for therapeutic applications.

## Methods

### Numerical solution of the molecular concentrations at the steady state

Consider a set of *K* chemical reactions involving *M* molecular species:

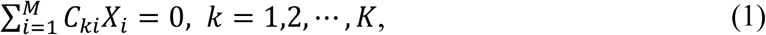

where *X*_*i*_ is the *i*’th molecular species and *C*_*ki*_ is the stoichiometry of *X*_*i*_ in the *k*’th reaction: *C*_*ki*_ > 0 indicates that *X*_*i*_ is a reactant, *C*_*ki*_ < 0 indicates that *X*_*i*_ is a product, and *C*_*ki*_ = 0 indicates that *X*_*i*_ is not involved in the reaction. The reactive flux for the *k*’th reaction is

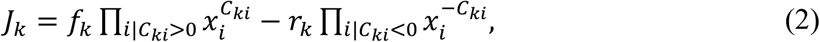

where *x*_*i*_ = [*X*_*i*_] is the concentration (or the activity in the case of non-ideal solutions) of *X*_*i*_ and *f*_*k*_ and *r*_*k*_ are the forward and reverse rate constants.

Let ***C*** be the *K*×*M* matrix of elements *C*_*ki*_ and ***J*** be the *K*-vector of components *j*_*k*_, the rate of change for the molecular concentrations is given by *d****x***/*dt* = −***C***^*t*^ · ***J***, with the superscript *t* denoting matrix transpose.

Let ***N*** = null(***C***) be the *M* × (*M* − rank(***C***)) matrix representing the null space of ***C***, then the time derivative of ***N***^*t*^ · ***x*** is

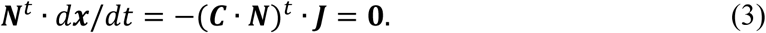

Thus

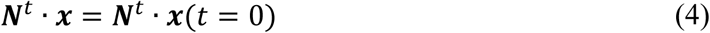

is constant, which reflects mass conservation; ***N***^*t*^ · ***x***(*t* = 0) gives the initial condition. At the steady state,

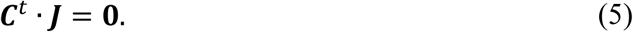

Equations 4 and 5 together constitute the steady state equations. To reduce them to linearly independent equations, take the singular value decomposition ***C*** = ***U*** · **Σ** · ***V***^*t*^. Let **Σ**_*s*_ be the rank(***C***) × rank(***C***) diagonal submatrix of **Σ** with the singular values being its diagonal elements and let ***U***_*s*_ be the *K* × rank(***C***) matrix consisting of the first rank(***C***) column vectors of ***U*** (these vectors span the range of ***C***). Equation 5 becomes

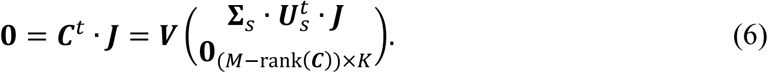

Because both ***V*** and **Σ**_*s*_ are full-rank, Equation 6 implies

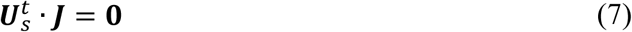

Equations 7 and 4 are the independent equations for the steady state concentrations.

### Checking that there is no spurious free energy production in the modeled set of reactions

Because my model is used to explore the conversion of ATP energy into non-equilibrium activation of Hsp90 clients, it must ensure that the total free energy change for any reaction cycle is zero. A reaction cycle is a subset of reactions that return all molecular species to their original states. It can be represented by a set of coefficients *Q*_*k*_ such that

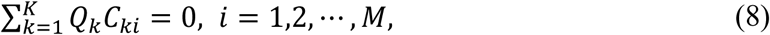

or in matrix form

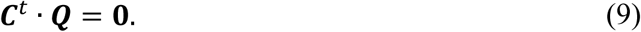

The reaction cycles thus correspond to the null space of *C*^*t*^. The total (unit-less) free energy change for any reaction cycle ***Q***,

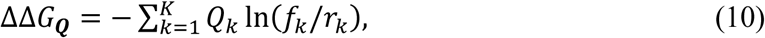

must be zero. This has been verified for the reactions in Tables 1 and 2 and the reaction of ATP hydrolysis in aqueous solution: ATP+H_2_O = ADP + Pi, which together constitute a closed chemical system with no external free energy input.

### Proof that a reaction can have a non-vanishing steady state flux only if it is part of a reaction cycle that consumes external free energy

Consider the set of reactions in Equation 1 at the steady state. The unit-less free energy change for the *k*’th reaction is

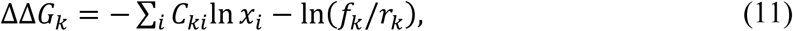

Equations 5 an9 imply that for any non-vanishing steady state flux ***J*** ≠ **0** we have a corresponding reaction cycle ***Q*** = ***J***. For any reaction cycle that does not consume external chemical energy, the total change of free energy must be 0, that is

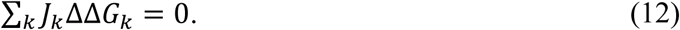

On the other hand, a reaction proceeds in the direction of free energy decrease, *i.e., J*_*k*_ΔΔ*G*_*k*_ ≤ 0, hence ∀*k, J*_*k*_ = 0. Q. E. D. To sustain ***J*** ≠ **0** at the steady state, chemical energy must enter the reaction cycle from molecules with an external source and sink.

## Author Contributions

HX confirms being the sole contributor of this work and this paper’s single author.

## Additional Information

The author declares no competing interests.

